# Deciphering the phospho-signature induced by hepatitis B virus in primary human hepatocytes

**DOI:** 10.1101/2024.04.10.588822

**Authors:** Florentin Pastor, Emilie Charles, Lucid Belmudes, Hélène Chabrolles, Marion Cescato, Michel Rivoire, Thomas Burger, Guillaume Passot, David Durantel, Julie Lucifora, Yohann Couté, Anna Salvetti

## Abstract

Phosphorylation is a major post-translation modification (PTM) of proteins which is finely tuned by the activity of several hundred kinases and phosphatases. It controls most if not all cellular pathways including anti-viral responses. Accordingly, viruses often induce important changes in the phosphorylation of host factors that can either promote or counteract viral replication. Among more than 500 kinases constituting the human kinome only few have been described as important for the Hepatitis B virus (HBV) infectious cycle, and most of them intervene during early or late infectious steps by phosphorylating the viral Core protein (HBc) protein. In addition, little is known on the consequences of HBV infection on the activity of cellular kinases.

The objective of this study was to investigate the global impact of HBV infection on the cellular phosphorylation landscape early after infection. For this, primary human hepatocytes (PHHs) were challenged or not with HBV, and a mass spectrometry (MS)-based quantitative phosphoproteomic analysis was conducted two- and seven-days post-infection. The results indicated that while, as expected, HBV infection only minimally modified the cell proteome, significant changes were observed in the phosphorylation state of several host proteins at both times points. Gene enrichment and ontology analyses of up- and down-phosphorylated proteins revealed common and distinct signatures induced by infection. In particular, HBV infection resulted in up-phosphorylation of proteins involved in DNA damage signaling and repair, RNA metabolism, in particular splicing, and cytoplasmic cell-signaling. Down-phosphorylated proteins were mostly involved in cell signaling and communication. Validation studies carried out on selected up-phosphorylated proteins, revealed that HBV infection induced a DNA damage response characterized by the appearance of 53BP1 foci, the inactivation of which by siRNA increased cccDNA levels. In addition, among up-phosphorylated RNA binding proteins (RBPs), SRRM2, a major scaffold of nuclear speckles behaved as an antiviral factor. In accordance with these findings, kinase prediction analysis indicated that HBV infection upregulates the activity of major kinases involved in DNA repair. These results strongly suggest that HBV infection triggers an intrinsic anti-viral response involving DNA repair factors and RBPs that contribute to reduce HBV replication in cell culture models.

## 1 INTRODUCTION

Despite the availability of an efficient preventive vaccine, nearly 300 million people worldwide are chronically infected with the Hepatitis B virus (HBV), making it a major contributor to liver diseases, especially cirrhosis and hepatocellular carcinoma (HCC) (https://www.who.int/news-room/fact-sheets/detail/hepatitis-b). Current antiviral treatments, primarily nucleotide analogs, can reduce viremia and the incidence of HCC (Wu et al., 2012). However, these treatments cannot completely eliminate the infection due to the persistence of a transcriptionally active viral genome in the nucleus of hepatocytes (Fanning et al., 2019).

The infection of human hepatocytes by HBV occurs through the recognition of its primary and secondary receptors on the cell surface, followed by the endocytosis of viral particles (Tsukuda and Watashi, 2020). After release into the cytosol, the capsid is directed towards the nucleus, where it binds to nuclear pores *via* its interaction with importins, before translocating through the nuclear channel and disassembling in the inner nuclear basket (Blondot et al., 2016;Diogo Dias et al., 2021). These early events result in the release of Core (HBc) protein, the unique structural component of the capsid, and the partially double-stranded (ds) circular genome known as relaxed circular DNA (rcDNA) to which the viral polymerase is covalently attached. The rcDNA is then repaired and loaded with cellular histones and other transcriptional regulators to form an extra-chromosomal covalently-closed circular DNA (cccDNA) which constitutes the template for production of viral RNAs (Diogo Dias et al., 2021;Wei and Ploss, 2021). During these early phases, two viral proteins also associate with cccDNA: HBc, derived from incoming capsids (240 monomers per capsid) or *de novo* translation (Lucifora et al., 2021;Locatelli et al., 2022), and HBx, the major HBV regulatory protein (Diogo Dias et al., 2021;Wang et al., 2023). Notably, HBx production by the infected cell is necessary for continuous viral transcription due to its capacity to induce the degradation of the smc5/6 complex (Lucifora et al., 2011;Decorsiere et al., 2016;Murphy et al., 2016;Niu et al., 2017;Abdul et al., 2022). The late phase of the HBV cycle occurs in the cytoplasm and involves the assembly of newly formed capsids in which the viral pregenomic RNA (pgRNA) is packaged along with the viral polymerase required for its reverse-transcription into rcDNA. Subsequently, these capsids are enveloped within multi-vesicular bodies and released into the extracellular space as infectious particles (Prange, 2022).

Most viral infections are characterized by the induction of strong intrinsic and innate anti-viral responses, against which viruses have developed sophisticated strategies to counteract them. In particular, viral infections can trigger a cellular DNA damage response (DDR), due to the recognition of viral genomes, which can prevent their replication (Weitzman and Fradet-Turcotte, 2018;Lopez et al., 2022). Cellular RBPs, participating in all steps of RNA metabolism, may also be engaged in a conflictual relationship with viral replication (Garcia-Moreno et al., 2018;Girardi et al., 2021;Lisy et al., 2021). In addition, cells can mount an innate response, intimately linked to the intrinsic one, resulting in the production of interferons or inflammatory cytokines which further amplify the anti-viral effect of both infected and neighboring cells (Guillemin et al., 2021;Justice and Cristea, 2022;Lopez et al., 2022).

In contrast to most viruses, infection of primary hepatocytes with HBV was reported to neither induce innate responses nor alter the transcription level of cellular genes and was, therefore, qualified as “stealth” (Mutz et al., 2018;Suslov et al., 2018). Nevertheless, whether HBV infection can tune cellular proteins by acting at a post-translational level, is presently unknown. In particular little is known about the effect of HBV infection on the activity of cellular kinases or phosphatases, and the downstream consequences on the phosphorylation of host proteins. So far, a unique study performed on infected hepatoma cells indicated that HBV can alter the host cell phospho-proteome and, in particular, target proteins involved in RNA processing, further suggesting that HBV infection could activate kinases involved in RBPs phosphorylation (Lim et al., 2022). Even though interesting, these results may be biased by the cellular model used for these analyses, HepG2 cells, which, as most cancerous cell lines, have dysregulated pathways, in particular those involved in viral sensing (Yang et al., 2014;Arzumanian et al., 2021).

The objective of this study was to investigate the global impact of HBV infection on the cellular phosphorylation landscape of PHHs. Our analyses indicate that HBV infection triggers important changes on the host cell phospho-proteome, in particular up-phosphorylation of several RBPs and some major factors implicated in DNA damage signaling and repair. Among these factors, we identified the RBP SRRM2, and the DNA damage sensor, 53BP1 as having antiviral activities that modulate the level of viral RNAs and cccDNA, respectively. Altogether, these analyses indicate that HBV infection is sensed by the host cell and triggers an anti-viral response mediated by changes in the level of phosphorylation of specific proteins that restrains viral replication.

## 2 MATERIAL AND METHODS

### 2.1 Cell culture and infection

HepaRG cells were cultured, differentiated, and infected by HBV as previously described (Gripon et al., 2002). PHHs were freshly prepared from human liver resection obtained from the Centre Léon Bérard and Hôpital Lyon Sud (Lyon) with French ministerial authorizations (AC 2013-1871, DC 2013 – 1870, AFNOR NF 96 900 sept 2011) as previously described (Lecluyse and Alexandre, 2010). HBV genotype D inoculum (subtype ayw) was prepared from HepAD38 (Ladner et al., 1997) cell supernatant by polyethylene-glycol-MW-8000 (PEG8000, SIGMA) precipitation (8% final) as previously described (Luangsay et al., 2015). The titer of endotoxin free viral stocks was determined by qPCR. Cells were infected overnight in a media supplemented with 4% final of PEG, as previously described (Gripon et al., 2002). Measure of secreted HBe and HBs antigens was performed by CLIA (Chemo-Luminescent Immune Assay) following manufacturer’s instructions (AutoBio, China) and expressed as international units/ml (IU/ml) and Paul Erlich Internation units/ml (PEIU/ml).

### 2.2 Cell extracts and mass spectrometry (MS)-based quantitative proteomic analyses

PHHs purified from liver resections were plated in 15-cm dishes (3 plates per time point) and 24 to 48 hours later infected with HBV (Multiplicity of infection (MOI) of 500 viral genome equivalents (vge)/cell) or mock-infected overnight. The following day, cells were washed with media then incubated until 2- or 7-days post-infection (dpi). Two independent experiments were performed: one with PHHs from one donor (PHH#1, 3 biological replicates per condition were prepared), and the other with PHHs from four donors (PHH#4). To prepare protein extracts, the cells were washed once with cold 1XPBS and then directly lysed on the plate using a buffer containing 8M Urea in 50mM Tris-HCl pH 8.0/75 mM NaCl/1mM EDTA supplemented with protease and phosphatase inhibitors (SIGMA). Cell lysates wee sonicated and frozen at −80°C.

Extracted proteins were reduced using 20mM of dithiothreitol for 1h at 37°C before alkylation with 55mM of iodoacetamide for 1h at 37°C in the dark. Samples were then diluted to ½ using ammonium bicarbonate and digested with LysC (Promega) at a ratio of 1:200 for 4 hours at 37 °C. Then they were diluted again to ¼ and digested overnight at 37 °C with sequencing grade-modified trypsin (Promega) at a ratio of 1:50. Resulting peptides were purified by C18 reverse phase chromatography (Sep-Pak C18, Waters) before drying down. Peptides were then labelled using an isobaric labelling-based approach, relying on tandem mass tags (TMT, (Thompson et al., 2003)) using the 16plex TMTpro isobaric Label Reagent kit (ThermoFisher Scientific) before mixing equivalent amounts and desalting using C18 reverse phase chromatography (Sep-Pak C18, Waters). An aliquot of labelled peptides was kept for total proteome analyses. Phosphopeptide enrichment was performed using titanium dioxide beads (TitanSphere, GL Sciences, Inc.) as previously described (Sonntag et al., 2017) before purification using C18 reverse phase chromatography (Marco SpinColumns, Harvard Apparatus). Isobaric-labelled peptides from total proteome and phosphoproteomes were then fractionated into eight fractions using the Pierce High pH Reversed-Phase Peptide Fractionation Kit (ThermoFisher Scientific) following the manufacturer’s instructions, except for the total proteome analysis of samples prepared from PHH#1 for which no fractionation was performed. The peptides were analyzed by online nanoliquid chromatography coupled to MS/MS (Ultimate 3000 RSLCnano and Q-Exactive HF, Thermo Fisher Scientific) using a 180 min gradient for fractions and a 480 min gradient if no fractionation was performed. For this purpose, the peptides were sampled on a precolumn (300 μm x 5 mm PepMap C18, Thermo Scientific) and separated in a 200 cm µPAC column (PharmaFluidics). The MS and MS/MS data were acquired by Xcalibur (version 2.9, Thermo Fisher Scientific). The mass spectrometry proteomics data have been deposited to the ProteomeXchange Consortium via the PRIDE (Perez-Riverol et al., 2022) partner repository with the dataset identifier PXD051216.

Peptides and proteins were identified and quantified using MaxQuant (version 1.6.0.17, (Tyanova et al., 2016)) searching in Uniprot databases (*Homo sapiens* and Hepatitis B virus taxonomies, 20210628 download) and in the database of frequently observed contaminants embedded in MaxQuant. Trypsin/P was chosen as the enzyme and two missed cleavages were allowed. Peptide modifications allowed during the search were: Carbamidomethyl (C, fixed), Acetyl (Protein N-term, variable), Oxidation (M, variable), and Phospho (STY, variable). Minimum peptide length and minimum number of razor+unique peptides were respectively set to seven amino acids and one. Maximum false discovery rates - calculated by employing a reverse database strategy - were set to 0.01 at peptide-spectrum match, protein and site levels.

Statistical analysis of quantitative data was performed using Prostar (Wieczorek et al., 2017). Peptides and proteins identified in the reverse and contaminant databases, and proteins only identified by site were discarded. Only class I phosphosites (localization probability ≥ 0.75) and proteins quantified in all replicates of at least one condition were further processed. After log2 transformation, extracted corrected reporter abundance values were normalized by the Variance Stabilizing Normalization (vsn) method, before missing value imputation (DetQuantile algorithm). Statistical testing was conducted with limma for results obtained with PHH#1 and two-tailed limma with paired design for results obtained with PHH#4, whereby differentially expressed proteins were selected using log2(Fold Change) and p-value cut-offs allowing to reach a false discovery rate inferior to 5% according to the Benjamini-Hochberg estimator. Proteins and phosphosites found differentially abundant but with imputed values in the condition in which they were found to be more abundant were manually invalidated (p-value = 1).

### 2.3 Bionformatic analyses

Gene ontology were performed using Metascape (Zhou et al., 2019) and the Reactome Gene Sets (http://reactome.org/), with a minimum overlap of 3, a minimum enrichment of 1.5, a p value of 0.01, and all genes as a background. Statistically enriched terms were identified, accumulative hypergeometric p-values and enrichment factors were calculated and used for filtering. Remaining significant terms were then hierarchically clustered into a tree based on Kappa-statistical similarities among their gene memberships. Then 0.3 kappa score was applied as the threshold to cast the tree into term clusters. Physical protein-protein interactions (PPIs) were similarly analyzed by Mescape with a minimum network size of 3. Protein networkformed by selected factors were also analyzed by Cytoscape (v3.10.1) with automatic network weighting. Prediction of kinases involved in the phosphorylation of differentially abundant phosphosites was performed using KinasePhos 3.0 (Ma et al., 2023).

### 2.4 siRNA transfection

dHepaRG cells or PHHs seeded into a 24-well plate were transfected with 25 nM of siRNA using Lipofectamine RNAiMax (Life Technologies), following manufacturer’s instructions. SiRNA used were the following: siHNRNPU (Dharmacon SmartPool L-03501-00), siSRRM2 (Dharmacon SmartPool L-015368-00), si53BP1 (Dharmacon SmartPool L-003548-00), siRIF1 (Dharamcon SmartPool L-027983) and siControl (Dharmacon D-001810).

### 2.5 Nucleic acid extractions and analysis

HBV RNAs and DNA were extracted from cells with the Nucleospin RNA (Macherey-Nagel) and MasterPureTM Complexe Purification Kit (BioSearch Technologies) kit without digestion with proteinase K (Allweiss et al., 2023), respectively, according to the manufacturer’s instruction. RNA reverse transcription was performed using Maxima First Strand cDNA Synthesis kit (Thermofisher). Quantitative PCR for HBV were performed using HBV specific primers and normalized to PRNP housekeeping gene as previously described (Lucifora et al., 2014). Pre-genomic RNA was quantified using the TaqMan Fast Advanced Master Mix (Life Technologies) and normalized to GusB cDNA levels. HBV cccDNA was quantified from total DNA by TaqMan qPCR analyses and normalized to β-globin cDNA level, as previously described (Werle-Lapostolle et al., 2004) or by droplet digital PCR using the “ddPCR Supermix for Probes (No dUTP)” (Bio-Rad) according to the manufacturer’s instruction. Droplets were generated using and the” QX200™ Droplet Generator” (Bio-Rad) and analyzed after PCR with the “QX600 Droplet Reader” (Bio-Rad).

### 2.6 Western blot analyses

Proteins were resolved by SDS-PAGE and then transferred onto a nitrocellulose membrane. Membranes were incubated with the primary antibodies corresponding to the indicated proteins. Proteins were revealed by chemi-luminescence (Super Signal West Dura Substrate, Pierce) using a secondary peroxydase-conjugated antibody (Dako) at a dilution of 1:10000. Primary antibodies used were anti-53BP1 (Abcam 175933, 1:1000), HNRNPU (Santa-Cruz sc-32315, 1:2000), β-tubulin (Abcam 6046,1:10000),

### 2.7 Immunofluorescence analyses

Analyses were performed as described previously using Alexa Fluor 555 secondary antibodies (Molecular Probes) (Salvetti et al., 2016). Primary antibodies used were: anti-53BP1 (Abcam 175933; 1:250); anti-PML Santa-Cruz sc-966, 1:250); anti HBc (Thermo MA1-7607, 1/500). Nuclei were stained with Hoescht 33258. Images were collected on a confocal NLO-LSM 880 microscope (Zeiss). Further image processing was performed using ICY (de Chaumont et al., 2012).

### 2.8 Statistical analysis

Statistical analyses were performed using the GraphPad Prism 9 software and a two-tailed Mann-Whitney non-parametric tests. A p value ≤ 0,05 was considered as significant. * correspond to p value ≤ 0.05; ** correspond to p value ≤ 0.01; *** correspond to p value ≤ 0.001.

## 3 RESULTS

### 3.1 Early impact of HBV infection on the host cell proteome and phosphoproteome

In order to examine the effect of HBV infection on the host cell phosphoproteome, we deployed a large-scale phosphoproteomic strategy. For this, HBV-infected PHHs, derived from a single donor, were lysed at 2- and 7-dpi (Figure 1A and B show experimental outline and levels of infection) before MS-based quantitative analysis of total proteome and phosphoproteome after phosphopeptide enrichment. The analysis of the total proteome indicated that the proteome remains largely unaffected by HBV infection, with less than 30 proteins found over- or under-expressed following infection at each time point, among the 3467 proteins identified and quantified (Figure 1C) (Supplemental Table 1). Interestingly, among proteins whose amount was increased at 2- and/or 7-dpi, figured, notably, fibronectin which was previously reported to be upregulated following HBV infection (Norton et al., 2004;Ren et al., 2016). In addition, two HBV under-expressed proteins at 7-dpi, LSM7 and TRIM21, were previously reported to negatively interfere with HBV replication (Song et al., 2021;Rahman et al., 2022). The deep phosphoproteomic profiling of these samples allowed to identify and quantify 8308 phosphopeptides containing class I phosphosites (localization probability > 75%) from 3012 different proteins (Supplementary Table 2). Among them, 161 and 316 were differentially abundant in PHHs infected by HBV compared to mock-infected PHHs, at 2- and 7-dpi, respectively (Fig. 1D) (Supplemental Table 2). A part of these modulated phosphosites, 17 up-regulated and 54 down-regulated in HBV-infected PHHs compared to mock-infected cells, were found differentially abundant at both time points. The comparison of total proteome and phosphoproteome results showed that only one protein, fibrinogen alpha chain (FIBA) but also its phosphorylated sites, were found to be more abundant in HBV-infected cells compared to control cells at 7-dpi; it was therefore excluded from further analyses since the measured up-regulation of its phosphosites could be linked to the over-expression of the protein. Overall, these results indicated that, in contrast to the minimal changes affecting the level of host proteins, HBV infection triggered more significant changes in the phosphoproteome, as early as 2-dpi, likely linked to the interplay between the virus and the host cell following attachment, internalization, and onset of viral replication.

**Figure 1.**
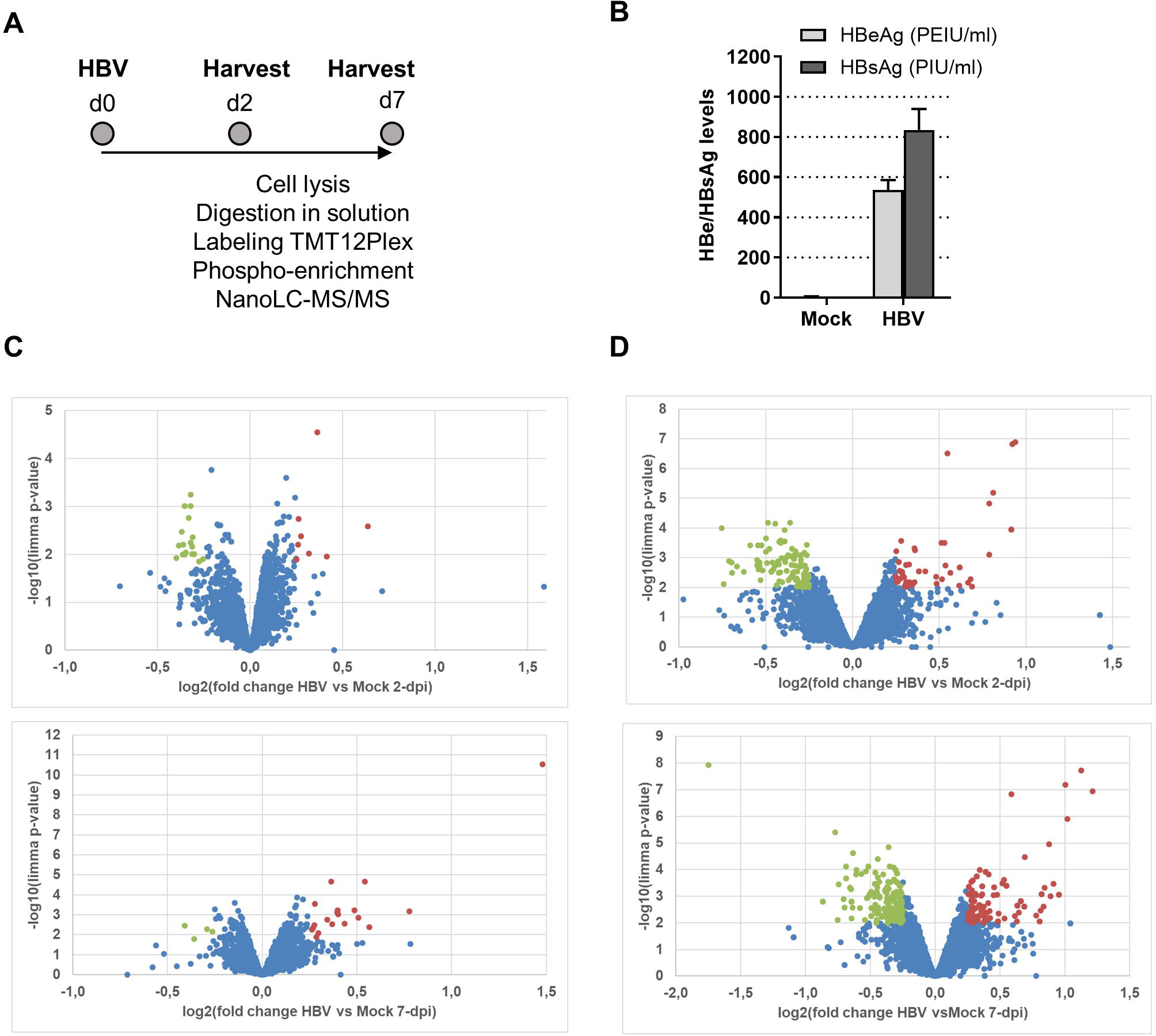
**A.** Outline of the experimental procedure. PHHs purified form liver resection were infected with HBV (MOI=500 vge/cell) overnight. Cells were kept in culture for 2 and 7 days, before direct on plate lysis and processing for MS and phospho-MS analyses. The experiment was performed in triplicate using PHHs from a unique donor. Mock: mock-infected cells. **B.** Quantification of HBe and HBs antigens levels in the supernatant of cells at 7-dpi. **C.** MS-based quantitative comparison of total proteomes of PHHs infected or not with HBV. **D.** MS-based quantitative comparison of phosphoproteomes of PHHs infected or not with HBV. Volcano plot display the differential abundance of proteins or phosphosites in cells infected or mock-infected with HBV for 2 (upper panel) and 7 (lower panel) days and analyzed by MS-based label-free quantitative proteomics. The volcano plots represent the -log10 (limma p-value) on y axis plotted against the log2(Fold Change HBV-infected vs mock infected) on x axis for each quantified phosphosite. Red and green dots represent, respectively, up- and down-phosphorylated proteins (left panels) and phosphosites (right panels) found significantly enriched in HBV-infected versus mock-infected samples at 2- and 7dpi (log2(Fold Change) ≥ 0.25 or ≤ −0.25 and p-value . 0.01, leading to a Benjamini-Hochberg FDR < 5%).

### 3.2 Analysis of host pathways modulated by HBV-induced up or down-phosphorylation events

In order to identify biological pathways and processes targeted by these phosphorylation events, gene ontology enrichment analyses were first performed on up-phosphorylated proteins (Figure 2A). This analysis indicated that HBV infection was characterized by up-phosphorylation of proteins involved in DNA repair pathways, notably non-homologous end-joining (NHEJ) and MAP kinase signaling, as early as 2-dpi. DNA repair factors were also targeted at 7-dpi, a time when other significant pathways appeared, notably those related to RNA splicing and cytoplasmic signaling by RhoGTPases. Analysis of physical protein-protein interactions (PPIs) based on human interactome datasets indicated that up-phosphorylated factors at both time points formed a network of proteins which contained densely-connected complexes related to double-stranded DNA breaks (DSBs), in particular NHEJ, and to MAPK-signaling (Figure 2B). A higher number of pathways were enriched when down-phosphorylated proteins were similarly analyzed, highlighting biological processes related to cytoplasmic signal transduction, in particular Rho-GTPases, which formed an important network of interconnected host factors (Figure 3A and B). Interestingly, EGFR, a host cofactor during HBV internalization (Iwamoto et al., 2019;Fukano et al., 2021), was found to be down-phosphorylated on serine 1166 and formed a major connected node at both time points (Figure 3B). Comparison of our data with those previously reported by Lim *et al*. (Lim et al., 2022), indicated that few up- and down-phosphorylated proteins were shared between the two studies (Supplemental Table 3). Many differences in the experimental set up may explain this low level of overlap, notably the use of primary (PHHs) versus transformed human hepatocytes (HepG2). Comparative analyses performed with factors previously described as interacting with HBc, and HBx also retrieved few common proteins (Chabrolles et al., 2020;Van Damme et al., 2021) (Supplemental Table 3). This latter observation suggests that most phosphorylation/dephosphorylation events observed in our study may not be associated to the capacity of these host factors to interact with viral proteins.

**Figure 2.**
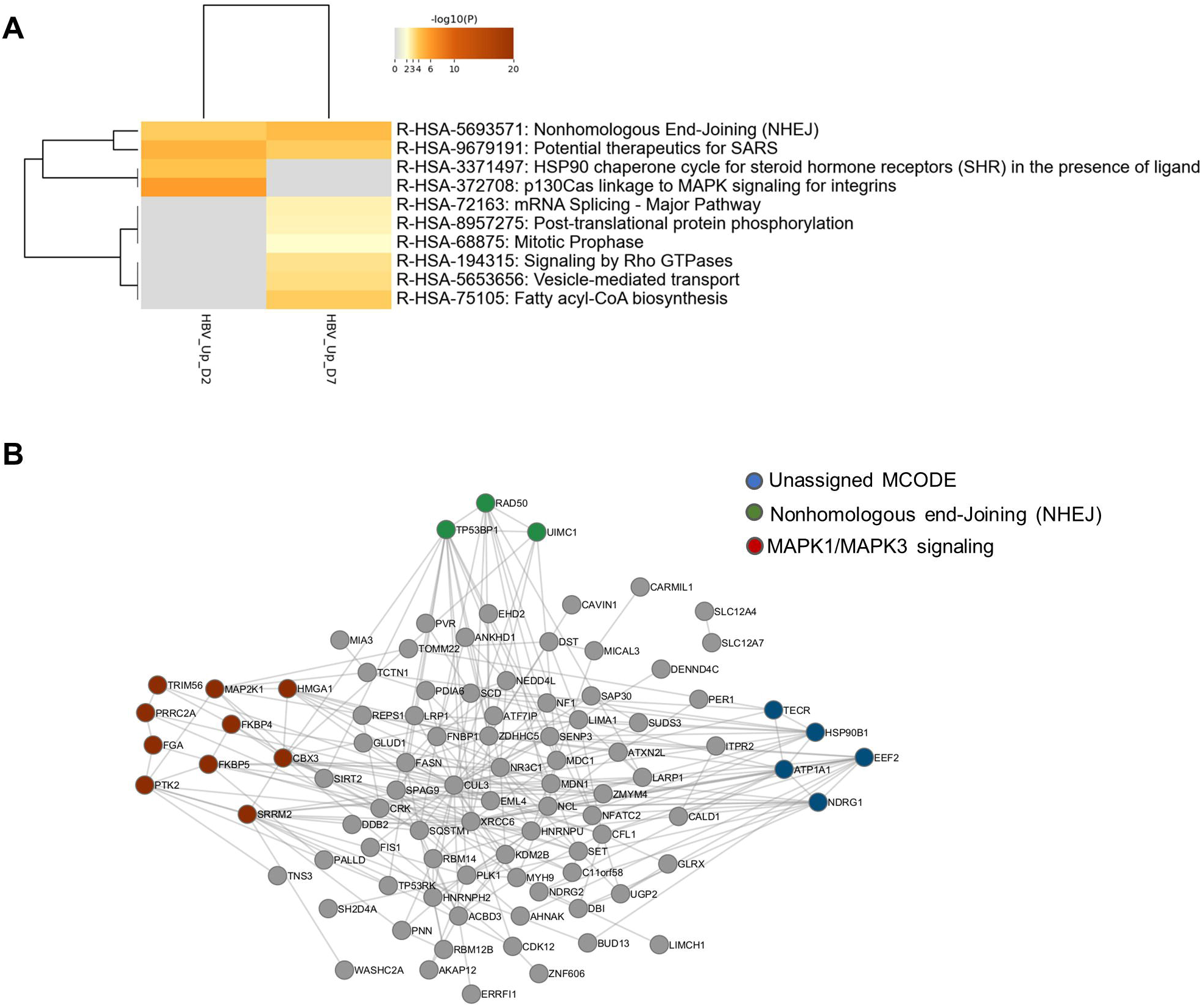
**A.** Gene ontology clusters formed by statistically enriched up-phosphorylated proteins at 2- and 7-pi. **B**. PPI networks formed by merged up-phosphorylated proteins at 2- and 7-dpi. All protein-protein interactions among input genes were extracted from PPI data sources using Metascape and a network MCODE algorithm was applied to identify neighborhoods where proteins are densely connected. Each MCODE network is assigned a unique color. Green: R-HSA-5693571|Nonhomologous End-Joining (NHEJ)|-5.1;R-HSA-5693565|Recruitment and ATM-mediated phosphorylation of repair and signaling proteins at DNA double strand breaks|-4.9;R-HSA-5693606|DNA Double Strand Break Response|-4.9; Red: R-HSA-8939211|ESR-mediated signaling|-4.3;R-HSA-5673001|RAF/MAP kinase cascade|-4.0;R-HSA-5684996|MAPK1/MAPK3 signaling|-4.0. Blue: unassigned. Grey: other interacting factors of the network.

**Figure 3.**
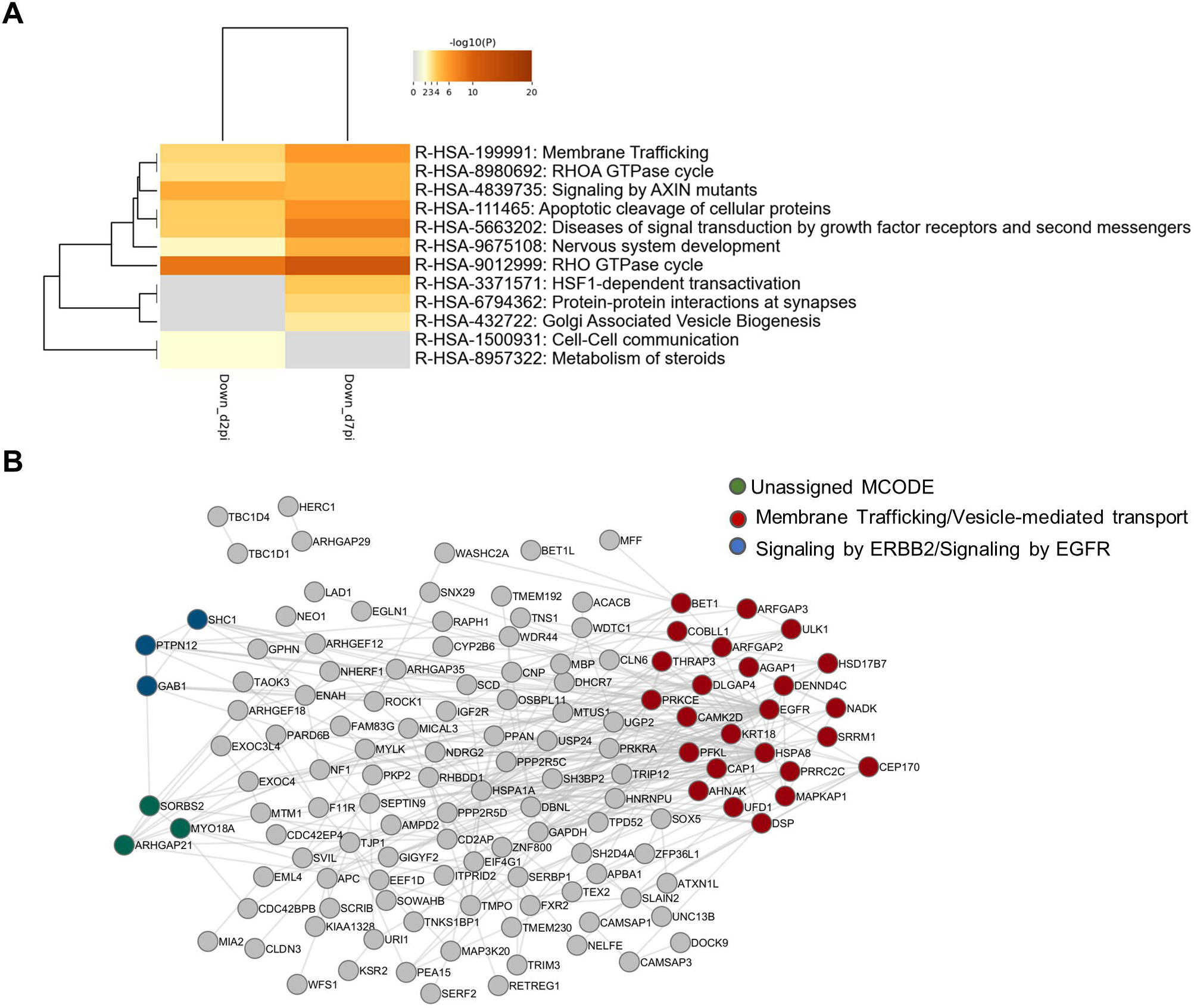
**A.** Gene ontology clusters of formed by statistically enriched down-phosphorylated proteins at 2- and 7-dpi. **B**. PPI networks formed by down-phosphorylated proteins at 2- and 7-dpi. All protein-protein interactions among input genes were extracted from PPI data sources using Metascape and a network MCODE algorithm was then applied to this network to identify neighborhoods where proteins are densely connected. Each MCODE network is assigned a unique color. MCODE annotation: Red: R-HSA-199991|Membrane Trafficking|-6.2;R-HSA-5653656|Vesicle-mediated transport|-6.1;R-HSA-6807878|COPI-mediated anterograde transport|-4.1. Blue: R-HSA-1227990|Signaling by ERBB2 in Cancer|-9.3;R-HSA-1227986|Signaling by ERBB2|-8.4;R-HSA-177929|Signaling by EGFR|-8.4. green: no annotation. Grey: other interacting factors of the network.

Altogether, these *in silico* analyses suggest that HBV infection triggers up- and down-phosphorylation events that target proteins involved in common pathways related to cytoplasmic signal transduction *via* Rho-GTPases as well as unique pathways, in particular related to DNA repair and RNA metabolism in the case of up-phosphorylated factors.

### 3.3 HNRNPU and SRRM2, two RBPs up-phosphorylated upon HBV infection, behave as anti-viral factor restricting HBV RNA production

To investigate the functional relevance of the results of our phospho-proteomic analysis, we first focused on up-phosphorylated RBPs since their activities are tightly regulated by phosphorylation, a post-translational modification which is frequently tuned by viral infections (Pastor et al., 2021;Velazquez-Cruz et al., 2021). In our analysis, several inter-connected RBPs were up-phosphorylated upon HBV infection. In particular, nine proteins (HNRNPH2, HNRNPU, SRRM2, PNN, NCL, BUD13, TP53RK, SET, and SENP13) constituted an enriched cluster of factors involved in mRNA splicing (Figure 4A). Of these, five RBPs, selected using the RBP2GO database (RBP2GO score above 40) (Caudron-Herger et al., 2021), formed an interconnected network (Figure 4B).

**Figure 4.**
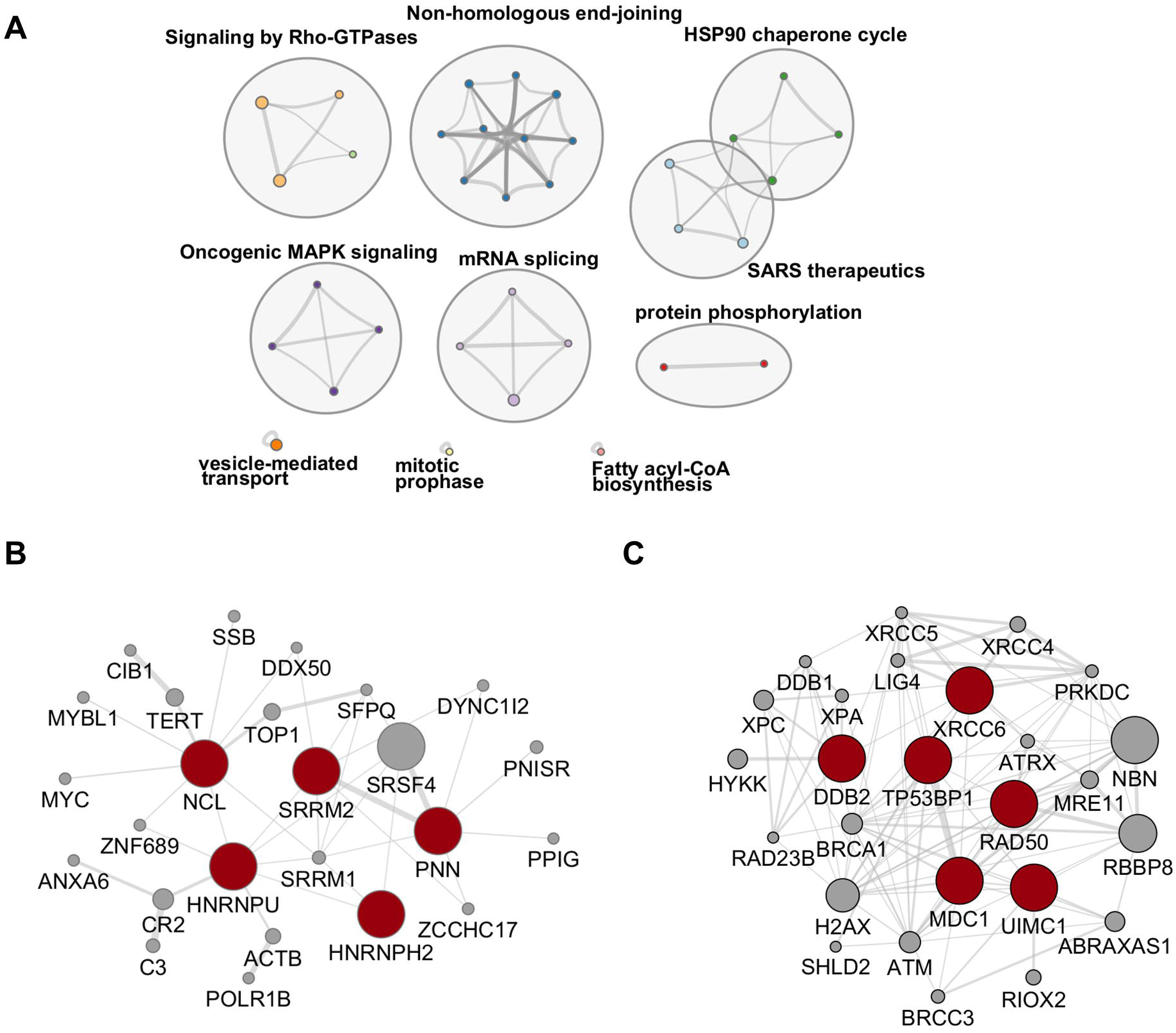
**A**. Network of enriched ontology clusters formed by up-phosphorylated proteins upon HBV infection. A subset of representative terms from the full cluster was converted into a network layout. Each term is represented by a circle node, where its size is proportional to the number of input genes that fall under that term, and its color represent its cluster identity (i.e., nodes of the same color belong to the same cluster). Terms with a similarity score > 0.3 are linked by an edge (the thickness of the edge represents the similarity score). The network is visualized with Cytoscape with “force-directed” layout and with edge bundled for clarity. **B**. and **C**. Protein-protein interaction networks formed by up-phosphorylated,proteins at 2- and 7-dpi involved in RNA binding (left panel), selected using the RBP2GO database (RBP2GO score above 40) (Caudron-Herger et al., 2021)and DNA repair (right panel), belonging to the NHEJ cluster. The physical interactions among up-phosphorylated proteins (red circles) were retrieved using by Cytoscape (v3.10.1) and the GeneMania application. Grey circles represent missing nodes used to build the interactome network.

Functional validation analyses were performed on two RBPs of this interconnected network: HNRNPU and SRRM2. HNRNPU, also called SAF-A, (for Scaffold Attachment Factor A), is a DNA- and RNA-binding protein that was identified as a constituent of the nuclear matrix capable of binding to nuclear matrix/scaffold-attachment regions (S/MAR) (Kiledjian and Dreyfuss, 1992;Romig et al., 1992;Jenke et al., 2002;Jenke et al., 2004). More recent studies indicate that HNRNPU is a chromatin scaffolding protein which, by oligomerizing with chromatin-associated RNA (caRNA), controls chromatin architecture and cellular gene expression (Nozawa et al., 2017;Fan et al., 2018;Xu et al., 2022). SRRM2, a member of the SR-related protein family, is involved in pre-mRNA maturation as a catalytic component of the spliceosome together with SRRM1, (Blencowe et al., 2000). Recent investigations also indicated that SRRM2, is a major component of nuclear speckles (Ilik et al., 2020;Xu et al., 2022). Both proteins, in particular SRRM2, are phosphorylated on multiple sites (http://www.phosphosite.org/). In this study, these proteins were found to be up-phosphorylated at a discrete number of sites, mainly on serine residues (Supplemental Table 2). To investigate the role of HNRNPU and SRRM2 during the HBV life cycle, knock-down (KD) experiments were performed on HBV-infected PHHs and differentiated HepaRG (dHepaRG) cells, using siRNAs (Figure 5). In both cell types, we found that knock-down of HNRNPU or SRRM2 resulted in an increase of HBV RNAs, both total RNAs and pgRNA, without significantly affecting cccDNA levels (Figure 5C and E). These results indicate that, as previously reported for other cellular RBPs, these factors behave as anti-viral factors that exert their functions at transcriptional and/or post-transcriptional level (Sun et al., 2017;Yao et al., 2018;Yao et al., 2019;Chabrolles et al., 2020;Yao et al., 2023b).

**Figure 5.**
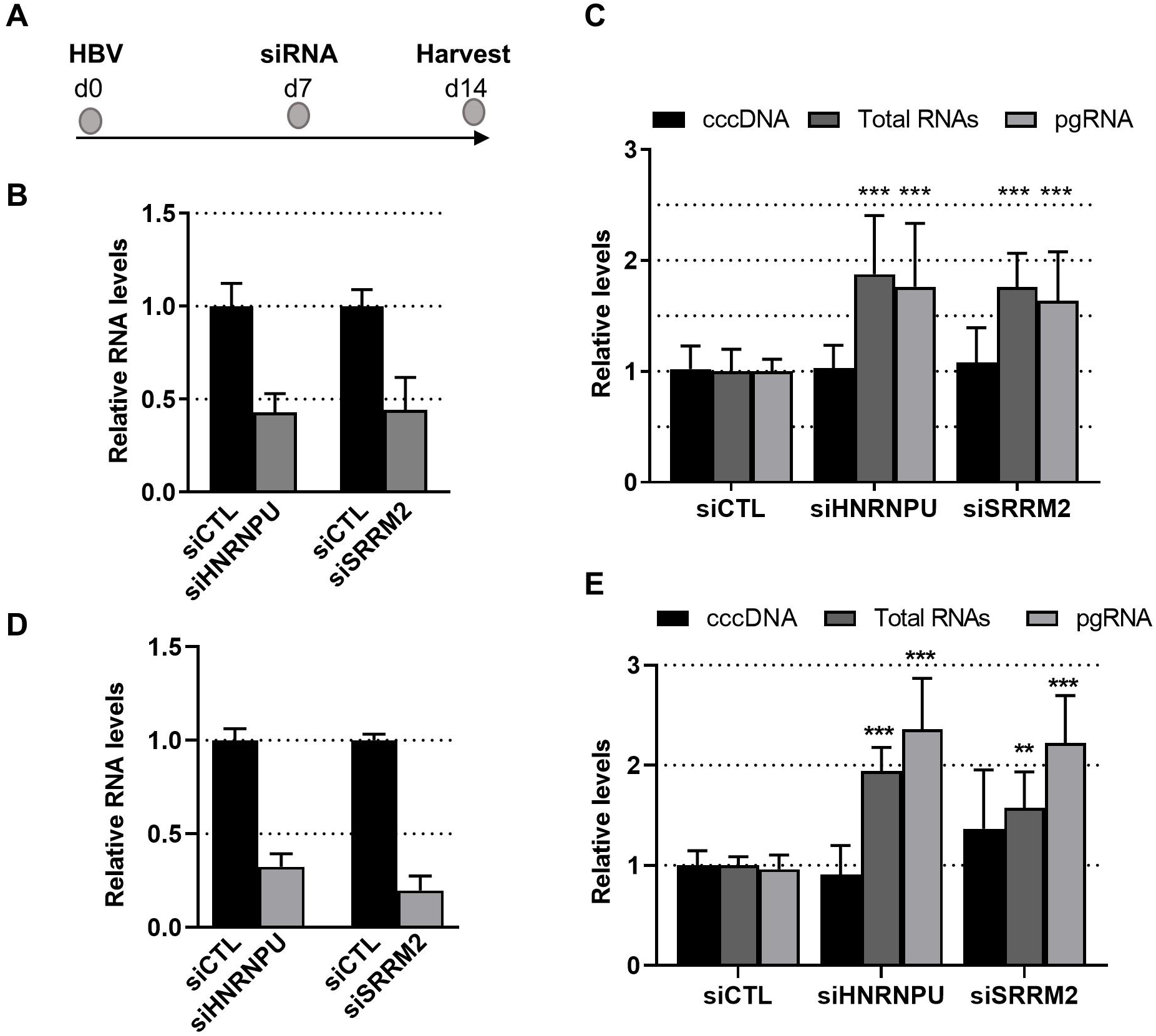
Effect of SRRM2 and HNRNPU knock-down on HBV life cycle. **A.** Experimental outline. PHHs (**B** and **C**) or dHepaRG (**D** and **E**) were infected with HBV (moi:500 vge/cell) and seven days later transfected with siRNA targeting HNRNPU, SRRM2 or control (CTL) siRNA. RNA and DNA were extracted from cells seven days after transfection to check the knock-down of cellular mRNA targets (**B** and **D**) and to quantify viral (**C** and **E**) nucleic acids. Results are expressed as the mean normalized ratio +/- SD, of at least 3 independent experiments, each performed in triplicate. Results in PHHs were obtained from four experiments performed with PHHs from four different donors.

### 3.4 HBV infection triggers an antiviral DNA damage response characterized by 53BP1 foci

DNA repair factors involved in NHEJ constituted another important cluster of proteins that were up-phosphorylated upon HBV infection (Figure 4A). Among these factors figured, notably, Ku70 (XRCC6), DDB2, Rad50, MDC1, and 53BP1 (Figure 4C). These proteins participate in a cascade of signaling events, including phosphorylation events, triggered by DNA damage, in particular by DSBs, that culminate with the recruitment of 53BP1 on chromatin, to form large foci that segregate damaged sites from the rest of the genome (Shibata and Jeggo, 2020;Rass et al., 2022). Even if 53BP1 foci formation can occur in the absence of phosphorylation following endogenous DSB produced during cell division, several reports have shown that stalled replication forks or exogenous genotoxic attacks induce the accumulation of phosphorylated 53BP1 at DSBs. (Anderson et al., 2001;Ward et al., 2003;Jowsey et al., 2007;Harding and Bristow, 2012). The resulting 53BP1 foci are essential for the assembly of DNA repair complexes and the initiation of the DNA damage response pathway (Shibata and Jeggo, 2020).

To investigate whether HBV infection could trigger the formation of 53BP1 foci, immunofluorescence (IF) analyses were performed on HBV-infected PHHs. We found that numerous 53BP1 foci were observed in HBV-infected cells, whereas only few foci were visible in mock-infected control cells (Figure 6A and B). Many of these foci, which appeared as early as 1-dpi, were located at the periphery of the nucleus. Importantly, these foci were observed using PHHs from different donors but were not observed, or only in few cells, when infection was performed in the presence of myrcludex, indicating that their formation may reflect a cell response to incoming HBV genomes (Figure 7). In addition, their presence was visible up to 7-dpi, when HBc staining was visible (Figure 7B). Altogether, these IF analyses strongly suggested that 53BP1 foci reflect a sustained host cell DNA damage response caused by incoming HBV genomes. In line with this hypothesis, we observed that in infected PHHs, most 53BP1 foci colocalized with PML bodies which were previously associated to long-lasting 53BP1 foci (Figure 8A and B) (Vancurova et al., 2019). In particular, 53BP1 appeared to form a scaffold around an inner PML core with some overlap at border (Figure 8C and D), strongly suggesting that these foci may constitute a hub of antiviral factors.

**Figure 6.**
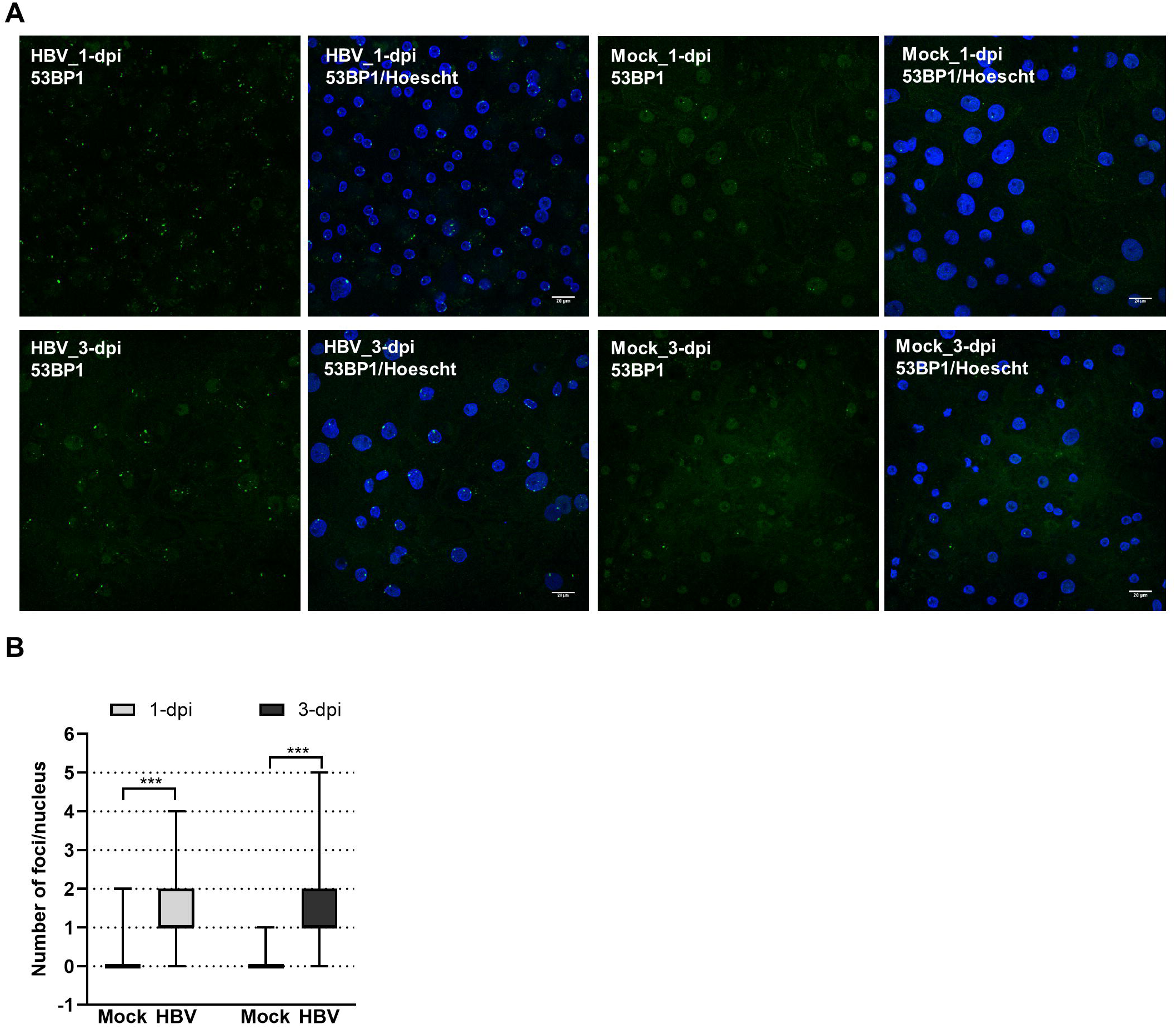
Visualization of 53BP1 foci in HBV-infected PHHs. **A.** HBV-infected (HBV) or mock (Mock)-infected PHHs were fixed at indicated time points and analyzed by immunofluorescence using an anti-53BP1 antibody (green signal), at 1- and 3-dpi. The nucleus was stained with Hoescht (blue signal). Scale bar: 20 µm. **B.** Box violin plot showing the number of 53BP1 foci per nucleus in HBV- or mock-infected PHHs, at both time points.

**Figure 7.**
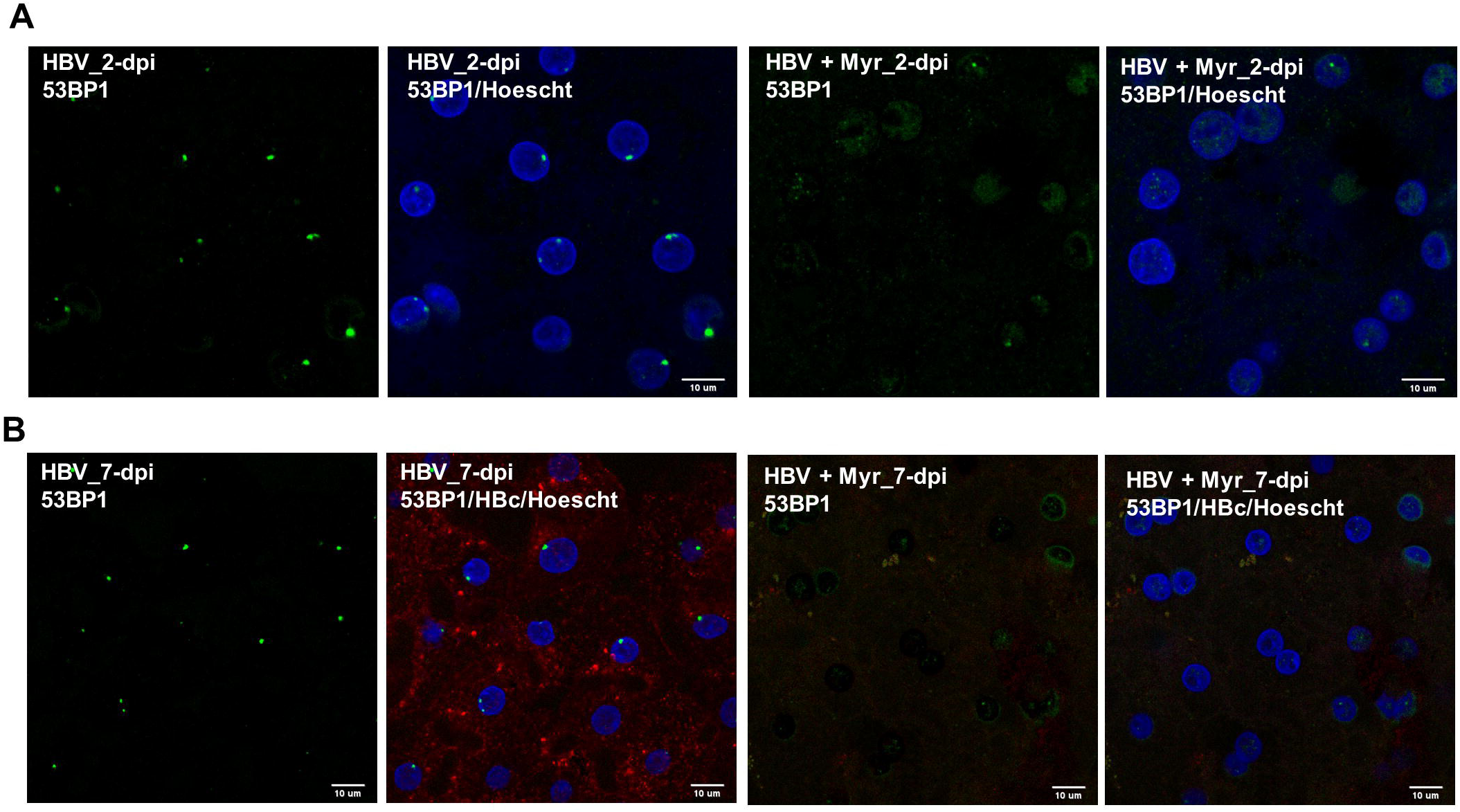
Co-labeling of 53BP1 foci in the presence of myrcludex. **A.** 53BP1 staining was performed on cells infected with HBV in the presence or not of myrcludex (Myr) at 2-dpi. Scale bar: 10 µm. **B.** HBV-infected PHHs, in the presence or not of myrcludex, were stained at 7-dpi using an anti-53BP1 (green signal) and an anti-HBc (red signal) antibody. Scale bar: 10 µm.

**Figure 8.**
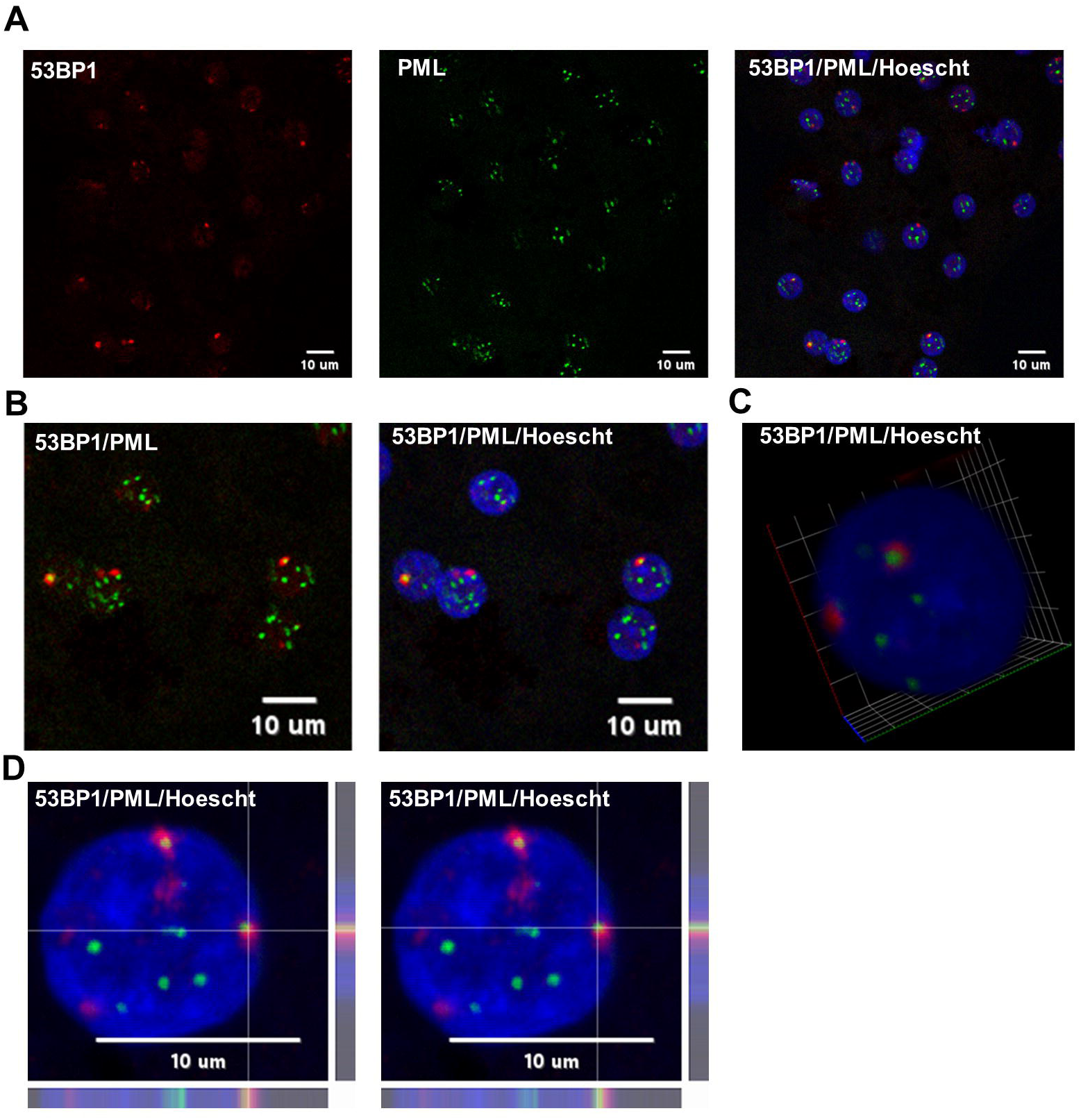
Co-labeling of 53BP1 and PML in HBV-infected PHHs. HBV-infected PHHs were fixed at 2-dpi and analyzed by immunofluorescence using an anti-53BP1 (red signal) and anti-PML (green signal) antibodies. The nucleus was stained with Hoescht (blue signal). Scale bar:10 µm. **B.** Enlarged merged views. **C.** 3D view of an hepatocytocyte nucleus showing the red (53BP1) signal surrounding the inner PML core (green signal). **D.** Line profile images of the same foci showing that the 53BP1 and PML signals partially overlap at the periphery of the inner PML core.

To determine whether 53BP1 may regulate the establishment of HBV infection, KD of this protein was performed before infecting hepatocytes with HBV (Figure 9A and B). Analyses performed 7 days later, indicated that depletion of 53BP1 resulted in a modest but significant increase in the level of cccDNA, strongly suggesting that this factor counteracts cccDNA establishment (Figure 9C). Recent studies have shown that upon binding to damaged DNA, 53BP1 recruits a shieldin complex which prevents long range resections required for DNA repair by homologous recombination (Setiaputra and Durocher, 2019). Recruitment of shieldin is mediated by RIF1 which recognizes phosphorylated residues on 53BP1 (Setiaputra et al., 2022). Interestingly, KD of RIF1 before HBV infection similarly increased the level of cccDNA and of viral RNAs, as observed for 53BP1 (Supplemental Figure 1).

**Figure 9.**
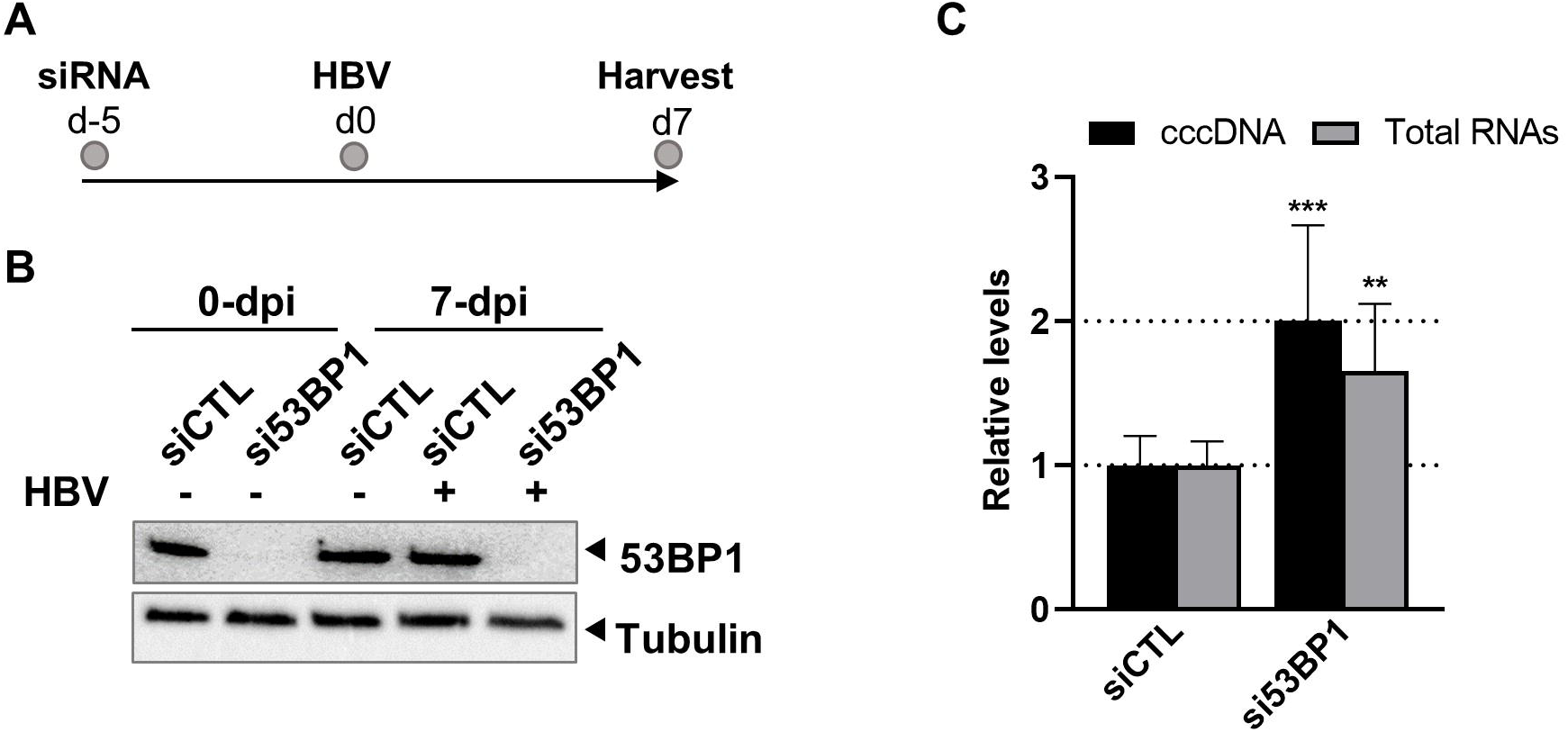
Effect of 53BP1 knock-down on HBV-infected dHepaRG. **A.** Experimental outline. **B.** Western blot validation of 53BP1 knock-down. **C.** Quantification of cccDNA and total HBV RNA levels. Results are expressed as the mean normalized ratio +/- SD, of three experiments, each performed in triplicate.

### 3.5 Extended phosphoproteomic analysis on PHHs derived from several donors and prediction of involved kinases

Our initial study of phosphosites modulated by HBV infection was performed on PHHs derived from a single donor. Therefore, an additional MS-based proteomic and phosphoproteomic analysis was performed on PHHs derived from 4 different donors. Strong donor-dependent variations in the efficiency of infection were observed among these four PHH batches (Supplemental Figure 2A). Not surprisingly, in these stringent conditions, of the 5731 proteins detected, very few were found to be differentially abundant in HBV-infected cells compared to mock-infected ones at 2- and 7-dpi (Supplemental Table 4 and Supplemental Figure 2B). Concerning phosphosites, 9179 were quantified, among which 54 and 60 were found significantly modulated at 2- and 7-dpi, respectively (Supplemental Table 5 and Supplemental Figure 2C). Several of them are from important proteins found to be differentially phosphorylated upon HBV infection in the previous analysis using PHHs from one donor, notably 53BP1 and SRRM2 (Supplemental Table 6). Enrichment analyses using the phosphoproteomic dataset obtained using PHHs from four donors indicated that HBV infection was characterized by up-phosphorylation of proteins involved in cell division and signal transduction (Figure 10A). PPI analysis indicated that most up-phosphorylated factors formed a connected network of proteins (Figure 10B). All the peptides found significantly up-phosphorylated in this latter experiment were analyzed with the KinasePhos3.0 software (Ma et al., 2023), to infer the most probable kinases activated by HBV infection at both time points. Interestingly, among the top most probable kinases involved in these modifications at both time points, figured, beyond the casein kinase II (CK2) which is implicated in a plethora of pathways (Borgo et al., 2021), ATM, ATR, and DNA-PK, the three major kinases activated upon DNA damage and which control the phosphorylation of downstream effector proteins (Figure11) (Blackford, 2017).

**Figure 10.**
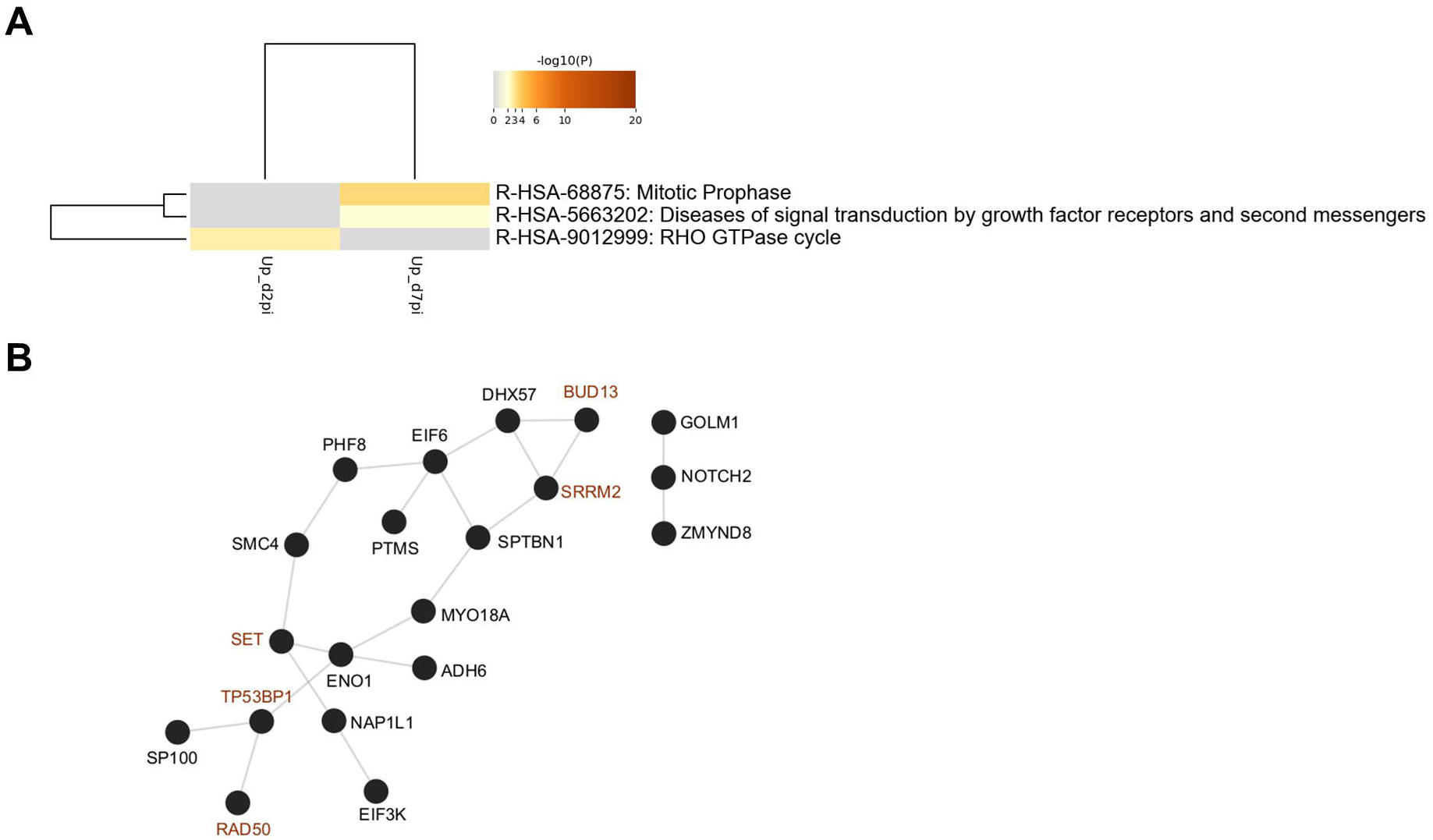
**A.** Ontology clusters formed by statistically enriched up-phosphorylated proteins at 2- and 7-dpi. (PHHs from four different donors). **B.** Physical interactions formed by proteins up-phosphorylated upon HBV infection. All protein-protein interactions among input genes were extracted from PPI data sources using Metascape. Red node names indicate proteins that were also found in the first phosphoproteomic analysis performed with PHHs from a single donor.

**Figure 11.**
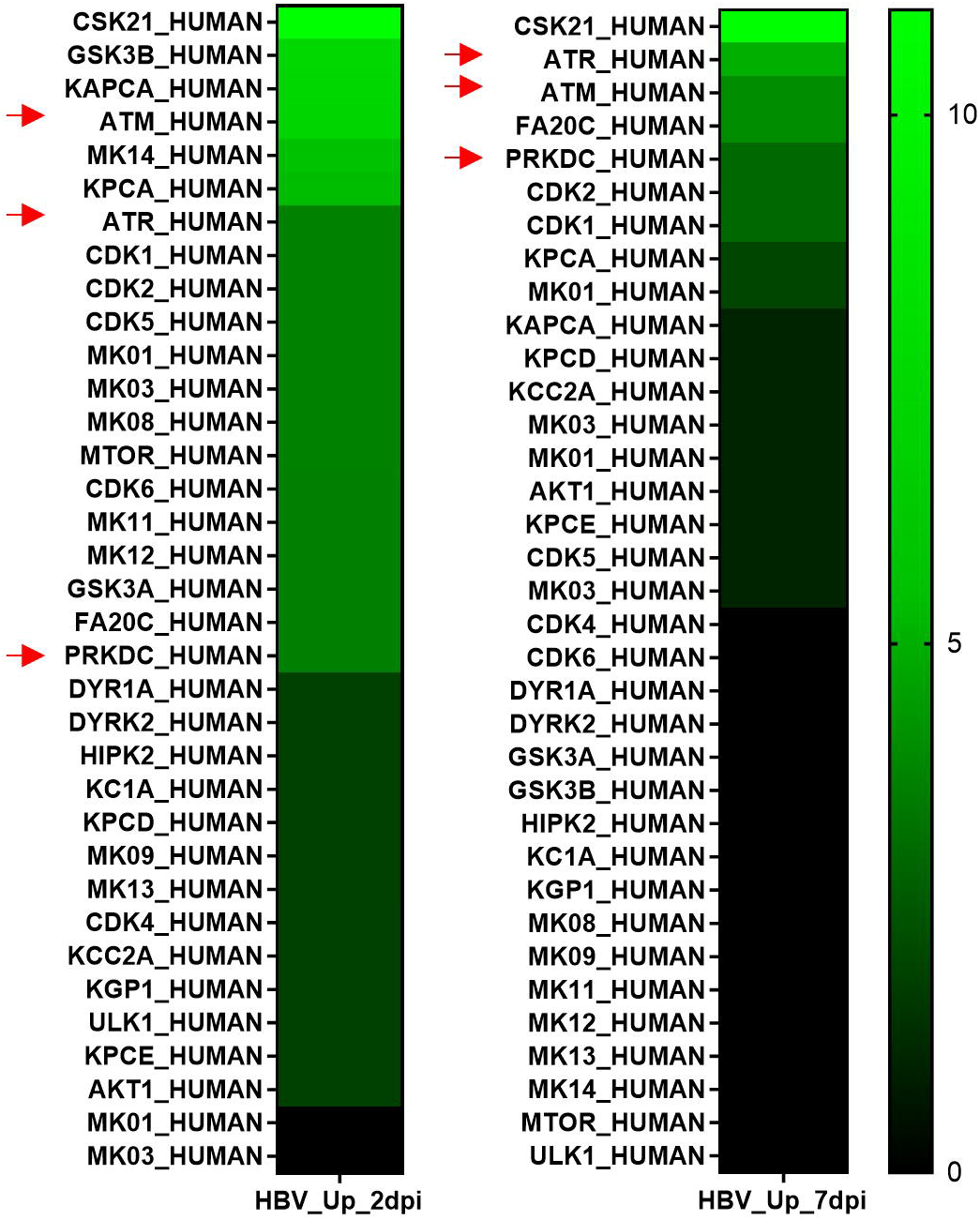
Cellular kinases predicted to be involved in up-phosphorylation events during HBV infection. Up-hosphorylated peptides found at d2- and d7pi were analyzed using KInasePhos3.0 software (Ma et al., 2023). The probability scores from 1 to 0.9, assigned to each kinase, were summed up to retrieve a list of the most probable kinases involved at each time point. Red arrows designate the three major kinases involved in DNA repair.

Altogether, this second study performed using PHHs from four different donors confirmed that HBV infection of PHHs induced the up-phosphorylation of a reduced but significant set of DNA repair proteins, in particular 53BP1. This finding correlated with the emergence of 53BP1 foci reflecting the induction of a DNA damage response. In addition, it also confirmed the up-phosphorylation of RBPs, such as SRRM2, a major constituent of nuclear speckles.

## 4 DISCUSSION

The conflicting interactions between a virus and the host cell during the early steps of the viral cycle are critical to determine the issue of the infectious process. In contrast to many other DNA and RNA viruses, infection by HBV was not reported to induce significant changes in host gene expression or innate responses, therefore leading to the assumption that HBV was able to enter the cell, and deposit its genome within the hepatocyte nucleus, without being detected (Mutz et al., 2018;Suslov et al., 2018). However, whether HBV infection may impact cell’s functions by acting at a post-translational level was still poorly documented. In particular, most viral infections can induce significant changes in the phosphorylation of host factors, reflecting altered kinase/phosphatase activities hijacked or induced upon viral entry, as shown for human imunodeficiency virus 1 (HIV-1) or severe acute respiratory syndrome coronavirus 2 (SARS-COV2), among others (Wojcechowskyj et al., 2013;Bouhaddou et al., 2020).

In this study we investigated whether HBV infection could alter the phosphorylation landscape of primary human hepatocytes which are growth-arrested and differentiated cells and, therefore, considered as the gold standard to study the HBV life cycle *in vitro* (Lucifora et al., 2020). We found that HBV infection can trigger both up-phosphorylation and down-phosphorylation of several cellular factors without significantly altering their expression level. As expected, many of these proteins were involved in pathways related to cell signaling. In particular, factors involved in signaling by MAPK were up-phosphorylated upon HBV infection. Even though signaling by these kinases are not essential for HBV infection, induction of this pathway may be linked to the requirement of EGFR for HBV entry (Iwamoto et al., 2019;Iwamoto et al., 2020). Interestingly, upon infection, EGFR was down-phosphorylated on serine 1166, whose activation by phosphorylation was previously reported to have negative impact on the EGFR activity (Assiddiq et al., 2012). In addition to MAPK-related pathways, factors involved in Rho GTPAses signaling were also detected among up- and down-phosphorylated proteins following infection, thus constituting an important signature. Rho GTPases control the actin cytoskeleton and, therefore, cell mobility, shape, and migration (Lawson and Ridley, 2018). Similarly, several effector proteins involved in the Rho GTPase signaling pathway and related to cytoskeletal organization, were also found in the phosphoproteomic analysis of SARS-CoV2-infected cells (Bouhaddou et al., 2020). A previous study, conducted by transfecting a plasmid containing the HBV genome in dividing HepG2 cells, reported that viral replication could induce morphological changes, activate Rac1 and, downstream, trigger the phosphorylation of ERK1 and AKT (Tan et al., 2008). Whether similar modifications can occur in more physiological infectious model is still unknown. In our analysis, neither AKT-nor ERK1-derived up-phosphorylated peptides were detected. Nevertheless, the finding that Rho GTPases signaling pathway was also highlighted in the more stringent analysis performed on PHHs from four different donors, strongly suggests that HBV infection may have an impact on the hepatocyte cytoskeleton and cell morphology.

Many studies documented the interaction between HBV and a plethora of RBPs that play a role at every step of viral RNAs production, from transcription to translation. In the present study, several RBPs were up-phosphorylated upon HBV infection. The HBc protein itself, which has a positively-charged, intrinsically disordered C-terminal domain similar to that found in many cellular RBPs, can also interact with a network of RBPs, some of which possess anti-viral activities (Diab et al., 2018;Chabrolles et al., 2020;Yao et al., 2023b;Zhang et al., 2023). Many if not all RBPs are tightly regulated by their phosphorylation level which controls their intra-cellular and intra-nuclear localization, their capacity to from condensates by liquid-liquid phase separation, their interaction with other protein partners, as well their affinity for RNA and DNA (Hofweber and Dormann, 2019;Velazquez-Cruz et al., 2021;He et al., 2023). In our study, we focused on two RBPs: HNRNPU and SRRM2. HNRNPU, is a DNA- and RNA-binding protein, which was recently described as chromatin scaffold factor able to regulate cellular gene expression, in particular by interacting with chromatin-associated non-coding RNAs (Sakaguchi et al., 2016;Nozawa et al., 2017;Fan et al., 2018). HNRNPU can also negatively regulate viral gene expression (Valente and Goff, 2006;Cao et al., 2019;Liu et al., 2021;Cao et al., 2022;Yang et al., 2022). Both up- and down-phosphorylated residues of HNRNPU were found in our study (Supplemental Table 2). In particular, serine 271 is described as a major target residue (https://www.phosphosite.org), whose phosphorylation is overrepresented during mitosis (Sharp et al., 2020). Interestingly, PLK1 which can phosphorylate HNRNPU and is activated upon infection (Douglas et al., 2015;Diab et al., 2017) was also up-phosphorylated in our first analysis performed with PHHs from a single-donor. As for HNRNPU, the absence of detectable up-phosphorylated peptides in the second analysis, performed with PHHs from four different donors, may be the consequence of strong variations among infections levels and/or donor-dependent features, in particular phosphorylation/dephosphorylation kinetics. The second selected RBP was SRRM2. Initially described as a component of the spliceosome and a member of the SR family of proteins, SRRM2 was recently identified as a main scaffold of nuclear speckles (Ilik et al., 2020). In particular, SRRM2 can form liquid condensates in a kinase-controlled fashion (Rai et al., 2018;Xu et al., 2022). This RBP was also up-phosphorylated in HIV-1-infected cells, in which it regulated alternative splicing of viral RNAs, as well as following infection of human macrophages with influenza A virus (Wojcechowskyj et al., 2013;Soderholm et al., 2016). Our validation studies indicate that the KD of either HNRNPU or SRRM2 is associated to a down regulation of HBV RNA levels without affecting cccDNA. However, as previously observed for SRSF10, an RBP which is part of the HBc interactome (Chabrolles et al., 2020), it is likely that the anti-viral effect of these proteins may vary according to their phosphorylation state. In particular, phosphorylation of speckle’s proteins, such as SC35 and SRRM2 by DYRK1A, DYRK3, and probably other kinases can dissolve liquid condensates formed by these speckle’s proteins, in particular during the onset of mitosis (Alvarez et al., 2003;Rai et al., 2018;Xu et al., 2022). Interestingly, in this study, DYRK1A figured among the kinases predicted to be activated at 2-dpi. Future investigation should focus on the ability of selected RBPs to interact with HBV cccDNA and/or RNAs in order to better understand the molecular mechanism involved in this antiviral effect. In addition, as for SRSF10, it will be important to investigate how HBV modulates the phosphorylation of these RBPs and, in particular, identify the cellular kinases involved.

The most remarkable finding of this study was the detection of up-phosphorylated proteins involved in DNA damage response and repair, strongly suggesting that these proteins could sense HBV infection, as previously shown for other nuclear viruses (Weitzman and Fradet-Turcotte, 2018). Notably, among these proteins, DDB2 was previously involved in cccDNA formation (Marchetti et al., 2022). In addition, some major DNA repair proteins, notably Rad50 and 53BP1 were found to be up-phosphorylated upon infection. In particular, Rad50, a cohesin-like component of the MRN complex, was up-phosphorylated at serine 635, a major ATM target (Kinoshita et al., 2009;Gatei et al., 2011). In the case of 53BP1, a highly phosphorylated protein (www.phosphosite.org), two different serine residues, previously reported to be up-phosphorylated upon DNA damage induced by ionizing or UV radiations, were detected in our analyses (Matsuoka et al., 2007;Boeing et al., 2016). 53BP1 is a critical regulator of the cellular response to DSBs (Panier and Boulton, 2014). When recruited to DSBs and phosphorylated, in particular by the ATM kinase, 53BP1 forms large foci which are a typical signature of an ongoing DDR (Anderson et al., 2001;Kilic et al., 2019;Shibata and Jeggo, 2020). In our study, detectable 53BP1 foci, most of them positioned at the nuclear periphery, were observed in HBV-infected PHHs, as early as 1-dpi, strongly suggesting that their formation was induced upon virus disassembly and rcDNA delivery at the inner face of the nuclear lamina (Rabe et al., 2009). Indeed, rcDNA harbors many features which represent danger signals for the cell, such as ssDNA breaks, ssDNA regions and a covalently attached polymerase (Schreiner and Nassal, 2017). Because most viral DNA genomes, possess unconventional features, their release within the nucleus, frequently results in a DDR (Weitzman and Fradet-Turcotte, 2018). In line with this hypothesis, we found that 53BP1 KD increased cccDNA levels, suggesting that this factor may prevent repair of rcDNA to produce a closed, double-stranded DNA episome. Interestingly, KD of RIF1, which recognizes phosphorylated 53BP1 and recruits the shieldin complex, also increased cccDNA levels, (Setiaputra et al., 2022). This last observation together with the predicted activation of DDR kinases strongly suggests that foci observed following HBV infection do contain phosphorylated 53BP1.

Accumulation of 53BP1 to DSBs and recruitment of the shieldin complex prevents long range resection of broken DNA ends required for DNA repair by homologous recombination (Setiaputra and Durocher, 2019;Callen et al., 2020). Our results therefore suggest that a certain level of resection of rcDNA may be required to generate cccDNA. In particular, exonucleases like Exo1 or DNA2-BLM, which are prevented to access DNA by the shieldin complex, may be involved (Setiaputra and Durocher, 2019). Alternatively, it is possible that the shieldin complex simply prevents the access of other repair factors, such as endonucleases, to rcDNA (Wei and Ploss, 2021). Several alternative pathways for rcDNA repair may, however, coexist, explaining why a certain level of cccDNA is produced under physiological conditions. In our analyses, 53BP1 foci were observed until 7-dpi, indicating a persistent DDR. Even if cccDNA formation is achieved in approximately 3 days in infected PHHs (Locatelli et al., 2022), a residual level of rcDNA may still be present and responsible for the presence of such foci. Alternatively, nuclear recycling of newly synthetized rcDNA in the nucleus may be responsible for the induction of this long-lasting DDR. A previous report indicated that PML bodies are recruited by 53BP1 to persistent DNA damage lesions (Vancurova et al., 2019). In our analyses, most of HBV-induced 53BP1 foci colocalized with PML signal. Interestingly, some studies indicated that, in the absence of HBx, PML bodies are important for HBV transcriptional silencing by recruiting inhibitory factors and viral genomes (Niu et al., 2017;Li et al., 2022;Yao et al., 2023a). It would be interesting to perform immuno-FISH analyses able to discriminate rc and cccDNA to determine whether PML/53BP1 signals also associate with viral DNA or whether they represent an alternative anti-viral nuclear structure.

Few information is available on the interplay between HBV and the DDR, in particular with its three major kinases, ATM, ATR and DNA-PK (Blackford and Jackson, 2017). Initial reports indicated that HBV infection may activate ATR and some of its downstream targets, therefore suggesting a possible recognition of rcDNA as a damaged molecule (Zhao et al., 2008a;Zhao et al., 2008b). Later on, both ATM and ATR were described as enhancing HBV replication (Kostyusheva et al., 2019), and the ATR/Check1 pathway shown to up-regulate cccDNA formation (Luo et al., 2020a). More recently, DNA-PK was described to increase HBV transcription (Fan et al., 2022). In our analyses, these three kinases were predicted to be activated upon HBV infection, as soon as 2- and up to 7-dpi. As already explored in some studies, this finding opens the perspective to investigate the effect of inhibitors of these kinases on the HBV life cycle and, in particular during the early phase, on cccDNA formation and transcription. Interestingly, Lubyova *et al*. reported that HBc may be phosphorylated by ATM following a genotoxic stress (Lubyova et al., 2021). It will be interesting to determine how kinase inhibitors targeting ATM or other DDR kinases modify the capacity of capsid-derived HBc to associate with cccDNA and how this affects downstream events. Finally, it is worth noting that, besides DDR kinases, CDK2, which plays a major role in HBc phosphorylation and is recruited and packaged within HBV capsids, was also on the top-ranked kinases activated upon infection (Ludgate et al., 2012;Luo et al., 2020b).

In conclusion, our deep MS-based phosphoproteomic analyses strongly suggest that HBV infection triggers an intrinsic anti-viral response composed by DNA repair factors and RBPs that contribute to reduce HBV establishment and productive replication (Figure 12). Future analyses conducted with other HBV genotypes, in particular genotype B and C that have a high genetic diversity and are responsible for infections in Asia and other parts of the world (Liu et al., 2018;Elizalde et al., 2021), will be important to further identify cellular pathways and proteins that either positively or negatively regulate HBV infection. Understanding how HBV may evade or counteract some of these responses will be critical to pinpoint factors able prevent cccDNA establishment and expression, as well to investigate further antiviral strategies based on the use of kinase inhibitors.

**Figure 12.**
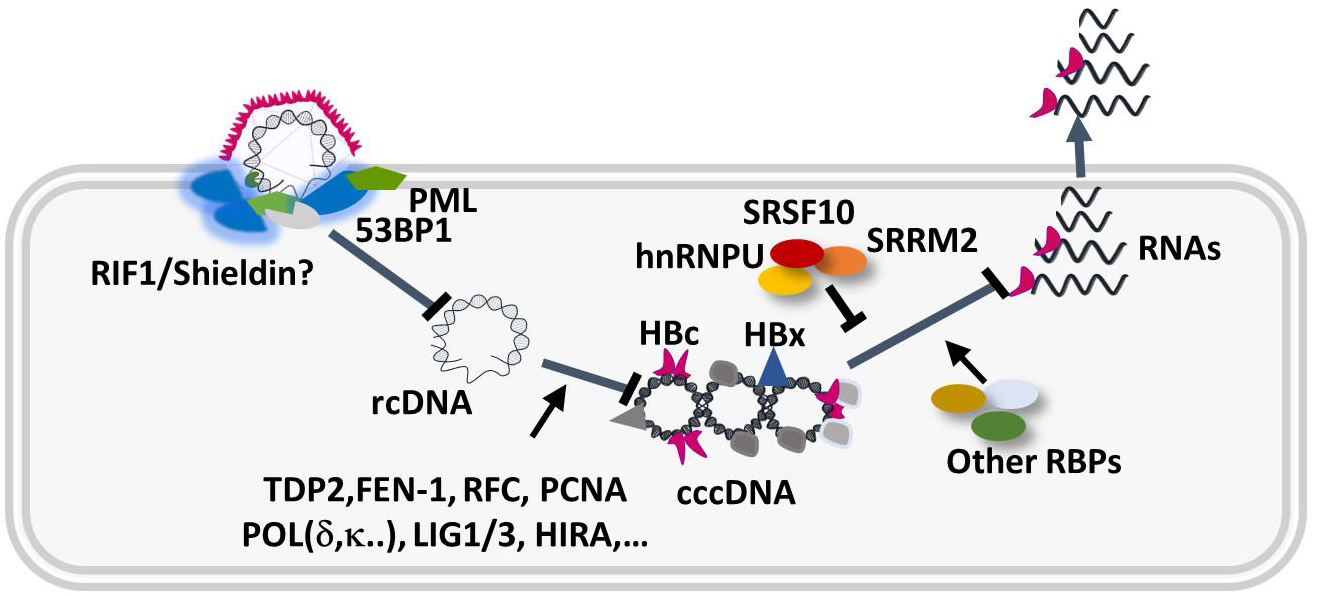
Hypothetical model describing the intervention of up-phosphorylated cellular DNA repair and RBPs during the HBV nuclear steps. Several host factors participate in cccDNA formation and viral gene expression (Diogo Dias et al., 2021;Wei and Ploss, 2021). Our study suggests that upon binding to the nuclear pore, and release of rcDNA at the inner side of the nuclear membrane, rcDNA is recognized as an abnormal molecule and induces a DDR characterized by ATM, ATR, activation and formation of 53BP1/PML foci. In addition, several RBPs, such as HNRNPU, SRRM2, and SRSF10 (Chabrolles et al., 2020), regulated by phosphorylation, further counteract HBV gene expression.

## 5 AUTHOR CONTRIBUTIONS

FP, EC, LB, HC and MC: Investigation, Methodology, Data curation. TB: Methodology. JL: Investigation, Writing – review & editing. DD: Funding acquisition, Writing – review & editing. MR and GP: Methodology. YC: Funding acquisition, Investigation, Methodology, Software: Data curation, Funding acquisition, Writing – review & editing. AS: Conceptualization, Funding acquisition, Project administration, Writing – original draft, Writing – review & editing.

## 6 FUNDING

This work was funded by Institut National de la Santé et de la Recherche Médicale (INSERM), the Centre National de la Recherche Scientifique (CNRS), and Université Claude Bernard Lyon 1 (UCBL). It was also supported by grants from the Agence Nationale de Recherche sur le Sida et les hépatites virales (ANRS, ECTZ11892) and fellowship to FP (ANRS, ECTZ119385). The proteomic experiments were partially supported by Agence Nationale de la Recherche under projects ProFI (Proteomics French Infrastructure, ANR-10-INBS-08) and GRAL, a program from the Chemistry Biology Health (CBH) Graduate School of University Grenoble Alpes (ANR-17-EURE-0003).

## 7 CONFLICT OF INTEREST

The authors declare that the research was conducted in the absence of any commercial or financial relationships that could be construed as a potential conflict of interest.

## 8 ACKNOWLEDGMENTS

We would like to thank Christophe Vanbelle (Imaging platform of CRCL), Ema Bobocioiu (PLATIM, imaging Platform of SFR Biosciences) for their help on confocal microscope analyses, Maud Michelet and Anëlle Dubois for their help for PHHs isolation, as well as the staff from Pr. Michel Rivoire’s and Dr Guillaume Passot’s surgery rooms for providing us with liver resections.

## List of abbreviations

caRNAs: chromatin-associated
RNAs cccDNA: covalently-closed circular
DNA DDR: DNA damage response
dGepaRG: differentiated
HepaRG dpi: days post-infection
DSBs: DNA double-stranded breaks
HBV: Hepatitis B Virus
HIV-1: human immunodeficiency virus 1
IF: ummunofluorescence
IU/ml: International Units/ml
KD: knock-down
MOI: multiplicity of infection
MS: mass spectrometry
NHEJ: non-homologous end-joining
PEIU/ml: Paul Erlich Institute Units/ml
pgRNA: pregenomic RNA
PHHs: primary human hepatocytes
PPIs: protein-protein interactions
RBPs: RNA-binding proteins
rcDNA: relaxed circular DNA
S/MAR: scaffold/matrix attachment region
SARS-Cov2: severe acute respiratory syndrome coronavirus 2
Vge/ml: viral genome equivalents/ml
Wt: wild type

## Notes

https://www.ebi.ac.uk/pride/login

### Competing Interest Statement

The authors have declared no competing interest.

### Summary of Updates

changes in the legend of Figure 1 change in Figure 1B Change in the legend of Supp Figure 2

