## Supplementary figures and images for "Deciphering the phospho-signature induced by hepatitis B virus in primary human hepatocytes"

### Supplemental Figure 1

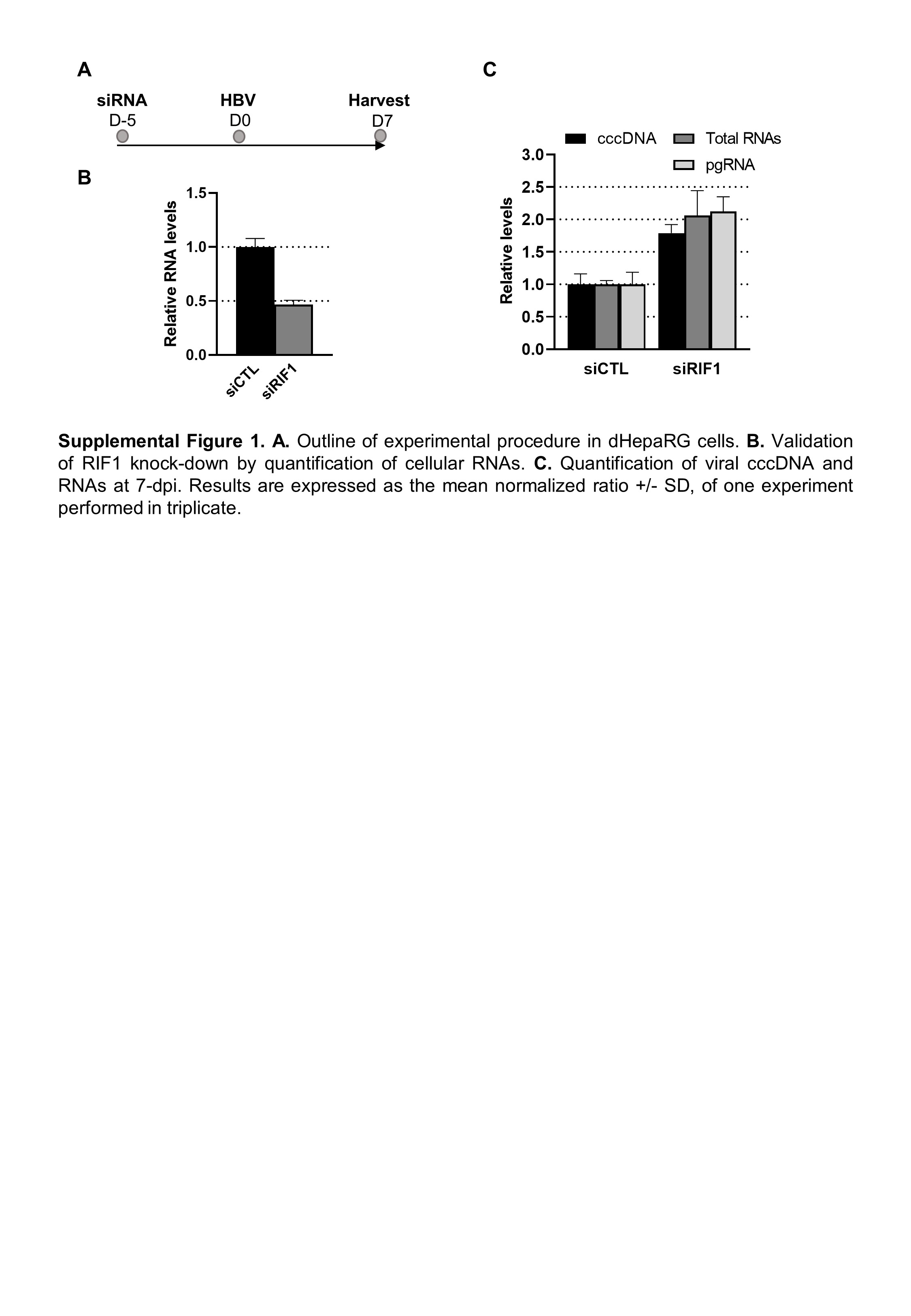

### Supplemental Figure 2

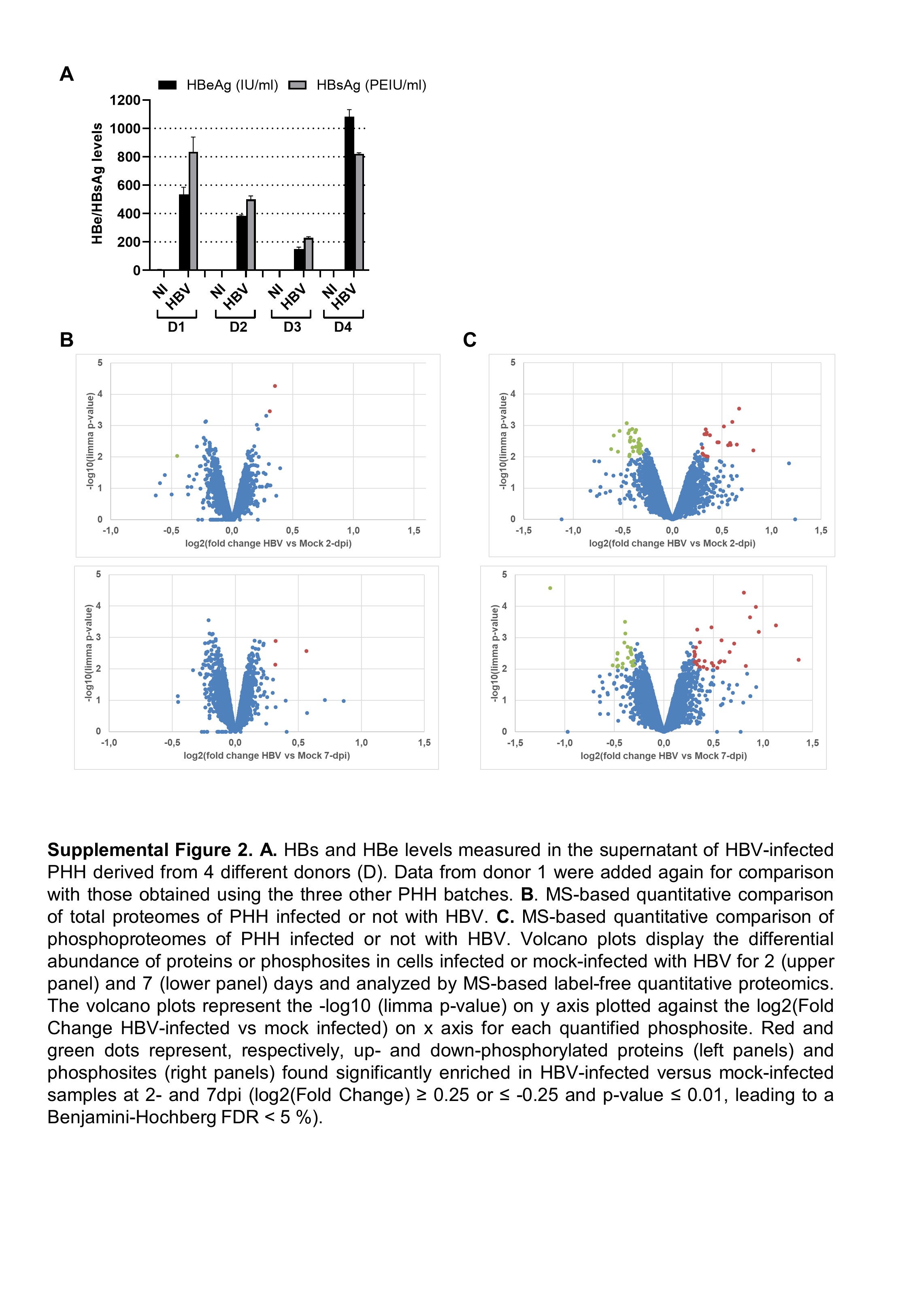
