## Supplemental Table 3 for "Deciphering the phospho-signature induced by hepatitis B virus in primary human hepatocytes"

|  | **This study** | | |
| --- | --- | --- | --- |
| **Method/targets** | **Up_d2pi** | **Up-d7pi** | **Up_d2&d7pi** |
| Phosphoproteomics  (Lim et al., 2022) | P13639 EEF2  Q9UQ35 SRRM2 | P48634 PRRC2A  P19338 NCL  Q9UJM3 ERBB  Q9H7L9 SDS3 | P49327 FASN  Q02952 AKAP12 |
| HBx  (Van Damme et al., 2021) | Q9Y4G6 TLN2  Q13469 NFAC2 | Q9Y3D6 FIS1  P12956 XRCC6 | - |
| HBc  (Chabrolles et al., 2020) | P13639 EEF2 | P23528 COF1 | - |
|  | **Down_d2pi** | **Down_d7pi** | **Down_d2&d7pi** |
| Phosphoproteomics  (Lim et al., 2022) | Q9Y2W1 TR150  E7EVA0 MAP4 | Q99959 PKP2  Q9UHD8 SEPTIN9  Q86WC4 OSTM1  A6ND36 FAM83G  P05783 KRT18  Q01433 AMPD2 | Q9Y5K6 CD2AP  P28290 ITPRID2  Q765P7 MTSS2  Q9C0C2 TNKS1BP1  Q8N6H7 ARFGAP2  P02686 MBP |
| HBx  (Van Damme et al., 2021) | - | P04406 G3P  P11142 HSP7C | P0DMV8 HS71A |
| HBc  (Chabrolles et al., 2020) | - | - | - |

**Supplemental Table 3.** Common cellular factors identified in this study and previous phosphoproteomics analyses or as interacting factors with viral HBx and HBc proteins. Each protein is designated by its Uniprot accession number and name.
