## Supplemental Table 6 for "Deciphering the phospho-signature induced by hepatitis B virus in primary human hepatocytes"

| **More abundant in**  **HBV-infected PHH** | **Less abundant in**  **HBV-infected PHH** |
| --- | --- |
| Q9BRD0 BUD13  P07108 DBI  QQ03001 DST  O43768 ENSA  Q8WX93 PALLDN  Q92878 RAD50  Q01105 SET  Q9UQ35 SRRM2  Q969E4 TCEAL3  Q9Y490 TLN1  Q12888 TP53BP1 | Q09666 AHNK  Q5XXA6 ANO1  Q13557 CAMK2D  Q13555 CAMK2G  Q9P1Y5 CAMSAP3  Q8WTX7 CAST  Q53SF7 COBLL1  O60716 CTNND1  Q9GZT9 EGLN1  Q13496 MTM1  Q15746 MYLK  O95544 NADK  Q9BXB4 OSBPL11  Q6ZUJ8 BCAP  Q70E73 RAPH1  Q13464 ROCK1  Q9H788 SH2D4A  P28290 SSFA2  Q07157 TJP1  P42166 TMPO  Q92890 UFDL1 |

**Supplemental Table 6.** Host proteins with modulated phosphosites upon HBV infection in both MS-based phosphoproteomic studies (if phosphosites are identical in the two studies, the gene names are underlined).
